# *Toxoplasma* does not secrete the GRA16 and GRA24 effectors beyond the parasitophorous vacuole membrane of tissue cysts

**DOI:** 10.1101/370502

**Authors:** Shruthi Krishnamurthy, Jeroen PJ Saeij

**Affiliations:** Department of Pathology, Microbiology and Immunology, School of Veterinary Medicine, University of California Davis, One Shields Ave., Davis, 95616, California, USA

**Keywords:** *Toxoplasma gondii_1_*, secreted effectors_2_, tissue cyst wall_3_, tetracycline inducible expression_4_, GRA16_5_, GRA24_6_

## Abstract

After invasion, *Toxoplasma* resides in a parasitophorous vacuole (PV) that is surrounded by the PV membrane (PVM). Once inside the PV, tachyzoites secrete dense granule proteins (GRAs) of which some, such as GRA16 and GRA24, are transported beyond the PVM likely *via* a putative translocon. However, once tachyzoites convert into bradyzoites within cysts, it is not known if secreted GRAs can traffic beyond the cyst wall membrane. We used the tetracycline inducible system to drive expression of HA epitope tagged GRA16 and GRA24 after inducing stage conversion and show that these proteins are not secreted beyond the cyst wall membrane.

## 1 Introduction

*Toxoplasma gondii,* which belongs to the phylum Apicomplexa, is an obligate intracellular parasite that can cause disease in immuno-compromised patients and fetuses (Montoya and Liesenfeld, 2004; Weiss and Dubey, 2009). It is the causative agent of toxoplasmosis, the 2^nd^ most common cause of food-borne illness in the USA (Jones and Dubey, 2012). Infectious tissue cysts are present in brain and muscles of many warm-blooded chronically infected hosts (Kim and Weiss, 2004). Infected cats, which are the definitive hosts, shed infectious oocysts in their feces contaminating food and water sources. Infection is initiated by ingestion of either tissue cysts containing the bradyzoite life-cycle stage or oocysts (Dubey, 1998). Upon stage conversion into tachyzoites and invasion of a host cell, *Toxoplasma* forms a parasitophorous vacuole (PV) that is surrounded by the PV membrane (PVM)(Black and Boothroyd, 2000). During invasion, *Toxoplasma* secretes effector proteins from rhoptries (ROPs), which mediate invasion, inhibition of host restriction factors, and modulation of host signaling pathways (Dubremetz, 2007; Boothroyd and Dubremetz, 2008). Once inside the PV, proteins from the dense granule secretory organelles (GRAs) are secreted onto, and beyond the PVM into the host cell cytoplasm (Hakimi and Bougdour, 2015). Three putative translocon proteins: Myc-regulation 1 (MYR1) along with MYR2 and MYR3 determine transport of GRA16 and GRA24 across the PVM into the host cell cytoplasm after which they traffic to the host cell nucleus (Franco et al., 2016; Marino et al., 2018). In addition to these putative translocon proteins, an aspartyl protease, ASP5 cleaves many secreted GRA proteins at a characteristic RRLxx motif also known as the *Toxoplasma* export element (TEXEL) motif which is important for their localization and function (Coffey et al., 2015; Hammoudi et al., 2015). Most of these effectors that have been characterized are from the non-orally infectious tachyzoite stage. It is unclear if bradyzoites within tissue cysts, akin to tachyzoites within the PV, can secrete GRAs beyond the PVM as the cyst wall is built on the inside of the PVM and presents a potential barrier for GRA secretion into the host cell.

GRA16, GRA24, GRA28 and IST (*T. gondii* inhibitor of STAT1 transcriptional activity) are secreted by tachyzoites beyond the PVM and traffic to the host nucleus (Bougdour et al., 2013; Braun et al., 2013; Gay et al., 2016; Nadipuram et al., 2016; Olias et al., 2016) where they modulate host signaling pathways important for parasite fitness. In this brief report we use the tetracycline inducible system to induce the expression of epitope tagged GRA16 and GRA24 after *in vitro* stage conversion. We observed that ATc induced *GRA16*-HA and *GRA24*-HA do not traffic to host cell nucleus. Instead, they accumulate within the *in vitro* tissue cysts.

## 2 Materials and Methods

### 2.1 Host cells and parasite strain

Human foreskin fibroblasts (HFFs) were used as host cells and were cultured under standard conditions using Dulbecco Modified Eagle Medium (DMEM) with 10% fetal bovine serum (FBS) (Rosowski et al., 2011). We chose GT1 parasites expressing Tet-R (Etheridge et al., 2014a), a type I strain that is capable of forming cysts during *in vitro* stage conversion (Lindsay et al., 1991; Fux et al., 2007) induced by pH 8-8.2+low CO_2_ (Skariah et al., 2010).

### 2.2 Plasmid construction

Using Gibson assembly (Gibson et al., 2009), we constructed Tet-On plasmids with a phleomycin resistance cassette to express GRA16-HA and GRA24-HA. The pTeton vector backbone was amplified with the following primers which was used to construct both the pTetONGRA16-HA SRS22A3’UTR as well as the pTetONGRA24-HA SRS22A3UTR constructs:

vector TetON for-GCATCCACTAGTGCTCTTCAAGGTTTTACATCCGTTGCCT
Vector TetON rev-AATTGCGCCATTTTGACGGTGACGAAGCCACCTGAGGAAGAC

The following primers were used to amplify the pieces for pTetONGRA16-HA SRS22A3’UTR Gibson assembly:

Vector GRA16 for-ACCGTCAAAATGGCGCAATTATGTATCGAAACCACTCAGGGATAC
SRS22UTRGRA16 rev-AATGACAGGTTCAAGCATAATCGGGAACGTCGTATG
GRA16HA SRS22A for-TTATGCTTGAACCTGTCATTTACCTCCAGTAAACATG
SRS22Avector rev-TGAAGAGCACTAGTGGATGCGTTCTAGTGCTGTACGGAAAAGCAAC

The following primers were used to amplify the pieces for pTetONGRA24-HA SRS22A3’UTR Gibson assembly:

vectorGRA24 for-ACCGTCAAAATGGCGCAATTATGCTCCAGATGGCACGATATACCG
SRS22AUTRGRA24HA rev-AATGACAGGTTTAAGCATAATCGGGAACGTCGTATG
GRA24HASRS22A For-TTATGCTTAAACCTGTCATTTACCTCCAGTAAACATG

### 2.3 Parasite transfection and selection

The pTetOn vectors containing GRA16-HA and GRA24-HA were linearized using AseI restriction enzyme. The linearized plasmids (50μg) were electroporated into 5x10^7^ GT1tetR parasites using the protocol described in (Gold et al., 2015). After lysis of parasites from host cells, they were selected twice with 50μg/ml phleomycin (Ble) and maintained in 5μg/ml Ble (Krishnamurthy et al., 2016) until they were cloned out by limiting dilution.

## Results

To test if GRAs are secreted beyond the tissue cyst wall membrane, we utilized the Tet-inducible system (Etheridge et al., 2014a) to express an HA-tagged copy of GRA16 or GRA24 under the Tet operator (Tet-O) in parasites expressing the tetracycline repressor (Tet-R). We could not just use *GRA16*-HA or *GRA24*-HA expressed from the endogenous promoter because if we see these proteins in the host nucleus we would not know if they were secreted beyond the PVM before the cyst wall was made (as tachyzoites) or after the cyst wall was made (as bradyzoites). In the absence of anhydrotetracycline (ATc), the Tet-R binds to Tet-O and represses the transcription of either *GRA16*-HA or *GRA24*-HA under the RPS13 promoter (Etheridge et al., 2014b; Wang et al., 2016). ATc binds TetR and relieves repression of transcription which allows for the expression of HA-tagged GRA16 or GRA24. Since ATc is smaller than the size-exclusion limit of the cyst wall (Lemgruber et al., 2011), we decided to use this system to answer our question.

To check if our constructs were able to stably express functional GRA16 and GRA24, we first transfected them into the RH parasite strain and observed nuclear localization of these proteins (data not shown). After transfection of the GT1 Tet-R expressing strain with the Tet-inducible GRA16-HA or GRA24-HA construct and subsequent selection with phleomycin for stable integration, we show by IFA that in tachyzoites GRA16-HA and GRA24-HA are only expressed in the presence of ATc (fig. 1a).

**Figure 1a.**
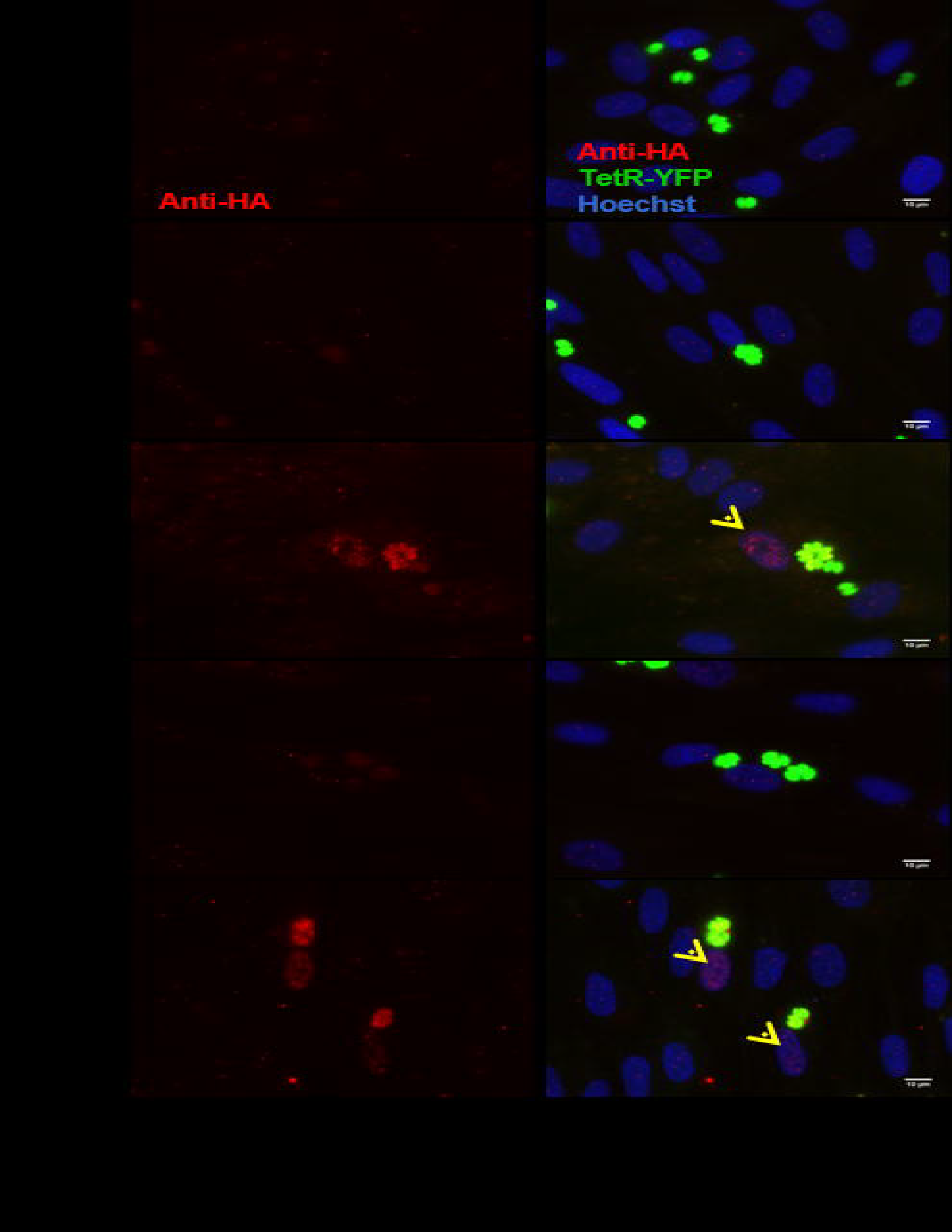
GT1-TetR parasites express GRA16-HA and GRA24-HA only in the presence of ATc. Coverslips with a monolayer of HFFs were infected in the presence or absence of 2μM ATC with GT1-TetR parasites or single clones of parasites expressing GRA16-HA and GRA24-HA under the RPS13 promoter downstream of a Tet-O7 operator. The coverslips were fixed 16 hours post infection with 3% formaldehyde and processed for IFA using rabbit anti-HA primary (Roche) antibody followed by goat anti-rabbit Alexa-555 secondary antibody and Hoechst (invitrogen). Tet-R is YFP tagged and GT1 tet-R parasites do not express any proteins that are HA-epitope tagged (first panel). In the absence of ATc, Tet-R represses the expression of GRA16-HA (second panel) and GRA24-HA (fourth panel). Repression by Tet-R is relieved only in the presence of 2μM ATc (third and fifth panel) allowing for the proper localization of GRA16-HA and GRA24-HA to the parasite PVM and host cell nucleus (yellow arrows). Images are scaled to 10μm.

A single parasite clone that expressed GRA16-HA or GRA24-HA only in the presence of ATc was chosen for induction of stage differentiation *in vitro* in human foreskin fibroblast (HFFs). The media was switched from DMEM with 10% FBS to tricine-buffered RPMI media with pH 8.0 and put in an incubator with low CO_2_ after 24 hours of infection (MOI=0.1) to induce tachyzoite to bradyzoite stage conversion. Five days post-switching, 2μM of ATc was added to the cultures to induce the expression of GRA16-HA and GRA24-HA since we observed that at least 50% of the parasites had converted to cysts by staining the cyst wall with DBA-lectin (Boothroyd et al., 1997) (data not shown). The parasites were fixed 48 hours following addition of ATc to allow for sufficient expression of GRA16-HA and GRA24-HA. We performed an indirect immunofluorescence assay (IFA) to determine the localization of GRA16 and GRA24 using anti-HA antibody as well as DBA-lectin to detect the cyst wall. We show that in host cells containing tissue cysts, GRA16-HA and GRA24-HA were not detected in the host cell nucleus or beyond the tissue cyst wall membrane and that instead they accumulated underneath the cyst wall. Almost 100% of vacuoles we observed were DBA positive. We decided to observe HFFs only infected with one parasite and therefore containing only one cyst as differences in the timing of conversion could affect the localization of GRA16 and GRA24. Out of 142 (59 for GRA16-HA and 83 for GRA24-HA from four biological replicates) images of singly infected host cells containing DBA positive cysts, both GRA16 and GRA24 were expressed exclusively beneath the cyst wall only in the presence of ATc (Figs 1b and 2a). We observed GRA24 localized to the host cell nucleus only in multiple infected cells containing tachyzoites, along with *in vitro* cysts (Fig. 2b). Thus, our results show that GRA16 and GRA24 are not secreted beyond the tissue cyst membrane into the host cell.

**Figure 1b.**
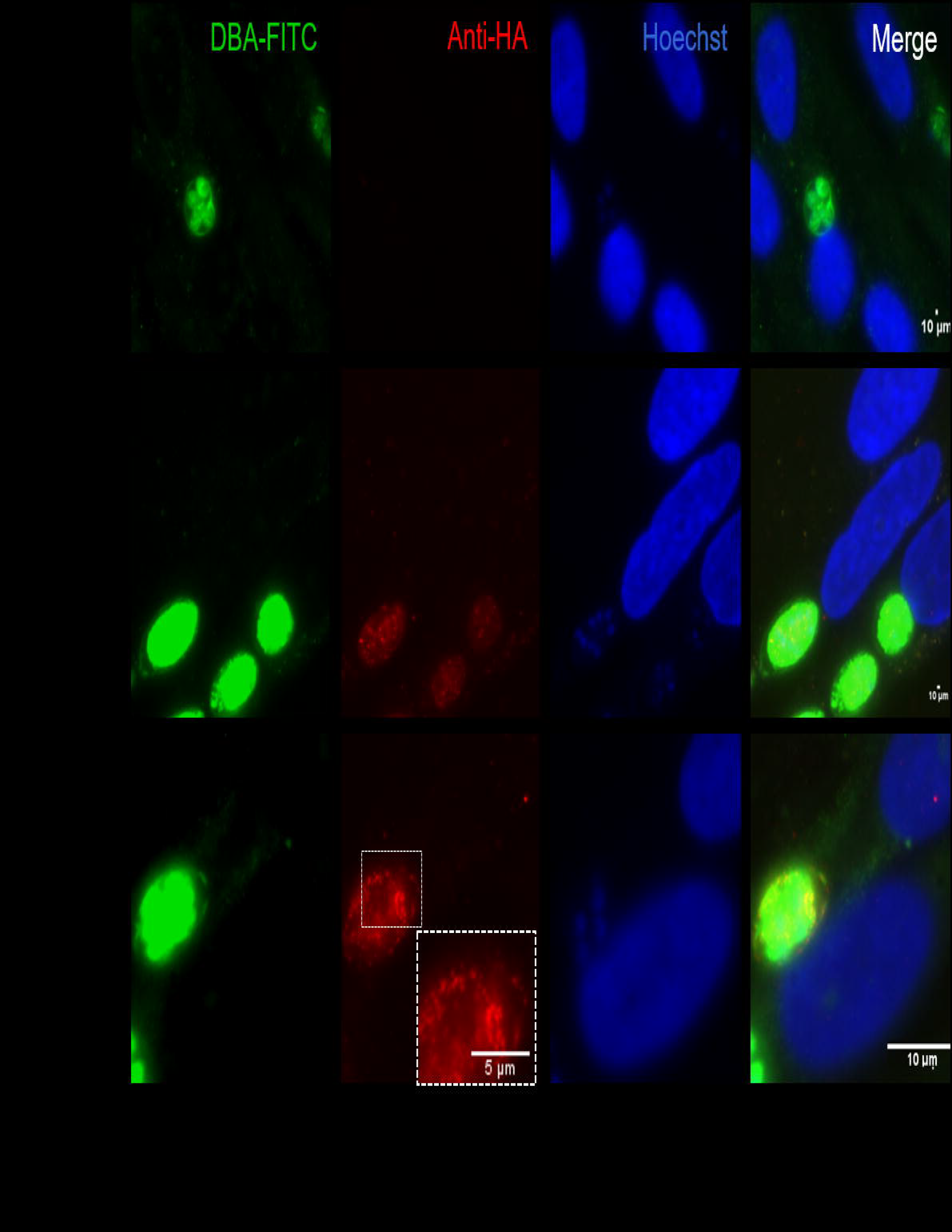
*In vitro* DBA positive cysts do not secrete GRA16-HA beyond the cyst wall membrane. HFFs were infected with a single clone of GT1-tetR parasites that expressed GRA16-HA only in the presence of ATc. After 24 hours of infection, stage differentiation was induced as indicated in the text. After 5 days, 2 μM ATc was added and 2 days later the coverslips were fixed with 100% cold methanol (also eliminates YFP signal from TetR) and processed for IFA. The cyst wall was stained with DBA-FITC along with HA and Hoechst. In the absence of ATc, GRA16-HA was not detected in parasites (first panel). GRA16-HA accumulated beneath the cyst wall only in the presence of ATc (second to fourth panels). Images are scaled to 10μm.

**Figure 2.**
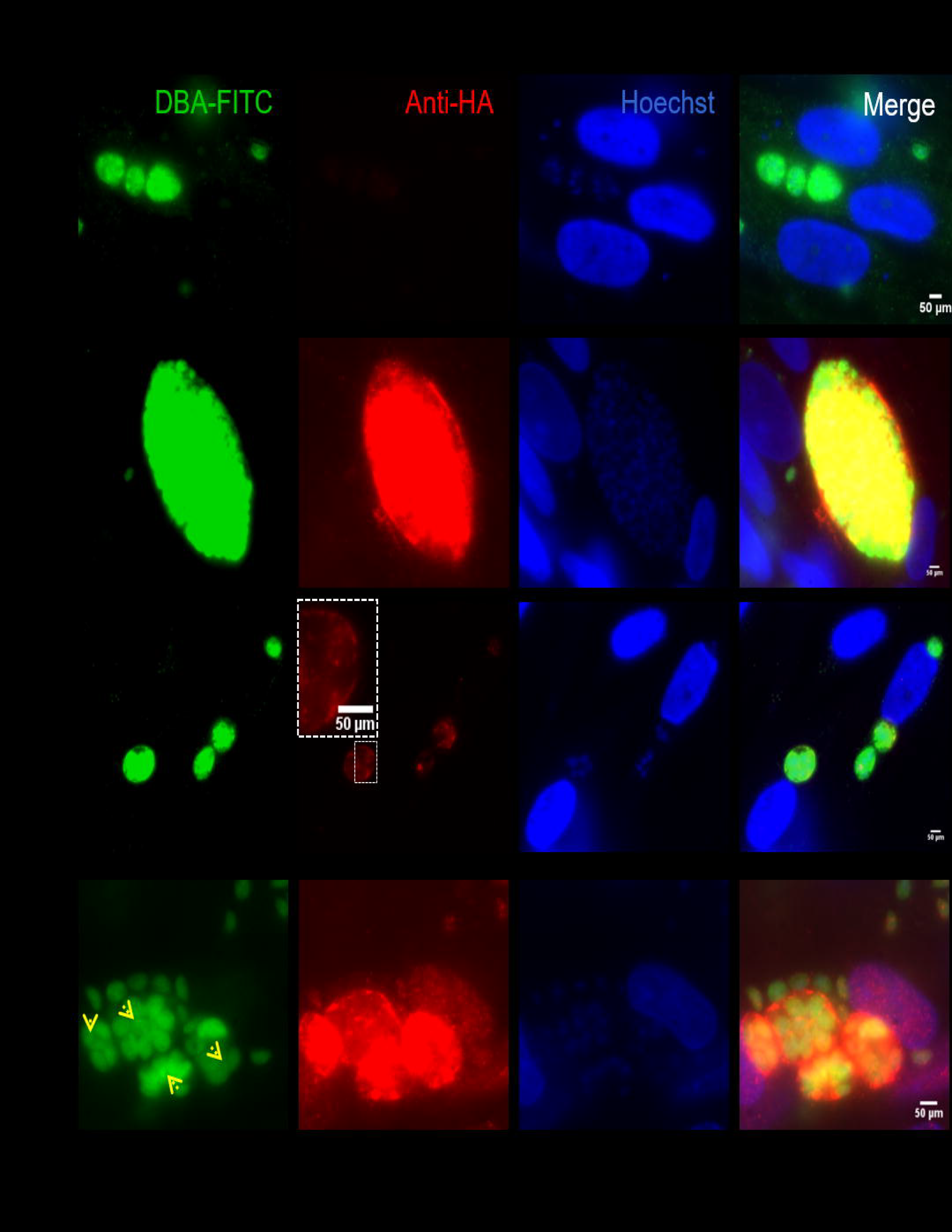
GRA24-HA accumulates underneath the DBA positive cyst wall. (a) Same IFA protocol was followed as described for figure 2. In the absence of ATc, there was no expression of GRA24-HA (first panel). In the presence of ATc, GRA24-HA accumulates beneath the cyst wall (second to fourth panel). (b) Multiple infected HFFs with tachyzoites and four *in vitro* cysts of parasites that express GRA24-HA showing nuclear localization. Yellow arrows indicate GRA24-HA expressing *in vitro* cysts. DBA-FITC staining is faint since 3% formaldehyde was used to fix parasites. Images are scaled to 50μm.

## 4 Discussion

We show here that GT1 parasites are able to form DBA lectin positive *in vitro* cysts. We also show for the first time that ATc is able to cross the cyst wall *in vitro*. Even though bradyzoites within tissue cysts are not as metabolically active compared to tachyzoites, it is becoming clear that they are also not in a dormant state (Sinai et al., 2016). However, we observed that bradyzoites do not secrete GRA16 and GRA24 beyond the *in vitro* cyst wall membrane. These proteins accumulated within the cyst wall suggesting that their role in the host cell nucleus is not required at this stage. Possibly bradyzoites require these proteins during natural oral infections after excystation from tissue cysts to establish infection in gut epithelial cells of the host. Not secreting parasite proteins beyond the cyst wall might help *Toxoplasma* to remain invisible and undetected by the host immune response during the chronic phase of infection. Another possibility may be that the translocon proteins MYR1/2/3 or ASP5 are not sufficiently expressed at this stage to effectively mediate transport of secreted GRAs beyond the cyst wall membrane. Even though all the MYRs and ASP5 are expressed in tachyzoites, sporozoites and bradyzoites, their expression is significantly lower in bradyzoites (Marino et al., 2018). However, even if MYR1-3 and ASP5 are expressed, our data indicate that the cyst wall seems to act as a barrier as ATc-induced GRA16 and GRA24 accumulated beneath the wall (Fig. 1 and 2).

## Abbreviations

Tet-on: tetracycline inducible system
TetR: tetracycline repressor protein
ATc: anhydrotetracycline
HA: hemagglutinin epitope tag
YFP: yellow fluorescent protein
Ble: phleomycin
HFF: human foreskin fibroblast
DBA lectin: dolichos biflorus agglutinin
IFA: indirect immunofluorescence
GRA: dense granule proteins
ROP: rhoptry proteins
IST: *T. gondii* inhibitor of STAT1 transcriptional activity
PV: parasitophorous vacuole
PVM: PV membrane

## Acknowledgments

We thank all the members of the Saeij lab for their feedback on this manuscript. We also thank Dr. Ronald D. Etheridge for proving us with GT1-TetR parasites.

## Author Contributions

SK generated all the data. SK and JS wrote and edited the manuscript.

